# Predictive Feature Engineering for Stress Detection using Physiological Signals, A Comparative Study

**DOI:** 10.64898/2026.07.24.740621

**Authors:** Jose Gabriel Gonzalez Nunez, Soheil Sabri, Parham Kebria, Jennifer Crook, Laura J Brattain

**Affiliations:** College of Engineering and Computer Science, University of Central Florida, Orlando, FL, USA; School of Modeling Simulation and Training, University of Central Florida, Orlando, FL, USA; Electrical and Computer Engineering, North Carolina A&T State University, Greensboro, NC, USA; College of Nursing, University of Central Florida, Orlando, FL, USA; College of Medicine, University of Central Florida, Orlando, FL, USA

**Keywords:** Electrodermal Activity, Foundation Models, Physiological Computing, Stress Detection, WESAD

## Abstract

This paper presents a two-stage pipeline for implicit feature engineering in time series-based physiological stress detection using electrodermal activity (EDA) signals. In the first stage, we forecast three descriptive statistics of future EDA signals over short horizons (3, 5, and 10 seconds) based on a 60-second context window. In the second stage, a lightweight linear classifier detects stress from these predicted statistics. We evaluate three forecasting architectures spanning the domain expertise spectrum: a domain-specific bidirectional long short-term memory (BiLSTM) recurrent neural network, zero-shot and fine-tuned variants of Amazon Chronos T5 time series foundation model, and the Tabular Prior-data Fitted Network (TabPFN) applied to engineered physiological features. Experiments on the publicly available Wearable Stress and Affect Detection (WESAD) dataset, comprising chest-worn multimodal physiological signals from 15 subjects under baseline and stress conditions, use subject-independent 5-fold cross-validation and show that the domain-specific BiLSTM and TabPFN achieve comparable classification performance, with mean area under the receiver operating characteristic curve (AUC) values of 0.859–0.882 and 0.863–0.883 respectively. Both remain well ahead of the Chronos variants, which yield 0.629–0.777. Chronos models quickly reach performance saturation regardless of training depth, highlighting challenges in tokenizing continuous physiological time series. The proposed approach advances implicit feature engineering for wearable stress monitoring by leveraging forecasting as a powerful inductive bias, thereby improving robustness and providing insights into the limitations of the foundation model for physiological signals.

## I. Introduction

Continuous monitoring of physiological stress is central to its application in the areas of occupational health, affective computing and digital twins of human performance models. Electrodermal activity (EDA) provides one of the most straightforward peripheral measures of arousal of the sympathetic nervous system, and hence is a commonly used modality in non-invasive stress detection systems [1]. EDA provides critical value to the development of human-centric Digital Twins (DTs) by serving as a high-fidelity, objective physiological marker for continuous stress monitoring and emotional state representation [2]. Within DT architectural frameworks, EDA data—often processed via bottom-up segmentation and physiological saliency cue (PSC) calculation—enables the creation of dynamic digital representations that accurately mirror and simulate individual distress responses to environmental or cognitive stimuli [3]. A significant innovation in DT development is the use of EDA features to train Gaussian Mixture Models (GMMs), which generate synthetic user profiles to augment training datasets, thereby enhancing the accuracy of stress detection classifiers such as Random Forest [2]. Furthermore, EDA acts as a vital temporal feature in deep learning pipelines, including CNN-LSTM and Temporal Convolutional Autoencoder (TCAE) models, which utilize compressed behavioral state vectors to predict personalized stress trajectories and detect early signs of mental health relapses [4]. Application areas for EDA-driven DTs are expansive: they are used in urban planning and Digital Twin City (DTC) models to map environmental stressors affecting older adults [3]; in User Experience (UX) evaluation to identify user discomfort during digital interactions [2]; in the educational sector to prevent student burnout through study-plan optimization [4]; and in clinical eHealth for the proactive management of cardiovascular abnormalities and chronic conditions like schizophrenia [5]. However, EDA is also inherently variable between and within individuals, making it challenging to extract stress-discriminative features.

To date, methods to extract such features can either be hand-crafted features (skin conductance level, response rate, amplitude, etc.) derived from windows of signal and then classified [6], or applied end-to-end learning architectures to raw signal [7]. Both paradigms process only observations and describe “what happened” rather than “what will happen”. Here, we instead focus on predicting future summary statistics of the EDA signal, and using these forecasts as features for classification.

There are two benefits to this prediction based approach. First, by predicting a summary statistic rather than the entire future signal (three numbers: mean change, variation and peak amplitude), we greatly reduce the dimensionality of the target prediction. Second, the process of forecasting is implicitly a feature engineering step; the model can only learn and predict parts of the future that are related to the observed past, effectively allowing the prediction itself to filter out uninterpretable physiological noise.

Given the recent advancements in foundation models for time series, an interesting question to ask is whether general-purpose pre-trained models are a good alternative to models that are specifically trained for physiological forecasting? We perform a comparison of three model families:

- A BiLSTM [8] with attention, trained from scratch on WESAD EDA data with five engineered input channels including cvxEDA-based tonic/phasic decomposition.
- Amazon Chronos [9], a time series foundation model built on the Text-to-Text Transfer Transformer (T5) architecture [10], evaluated in zero-shot mode and with fine-tuning of the T5 decoder (encoder-frozen).
- TabPFN [11], a tabular foundation model that performs incontext learning on engineered statistical features extracted from the lookback window, including tonic and phasic component statistics.

The primary contributions of this work are: (1) A new pipeline for generating features for EDA based stress detection via a learned prediction of near future signal summary statistics; (2) A subject-independent cross-validated comparison of domain-trained and foundation model architectures for physiological signal analysis; (3) An ablation study to examine the architectural shortcomings of foundation models that rely on tokenization for physiological signal processing; and (4) An empirical demonstration that learned sequential forecasting can yield features that are competitive with a near-perfect knowledge of the true future signal.

## II. Related Work

### A. Stress Detection from EDA

The Wearable Stress and Affect Detection (WESAD) dataset [6] is a publicly available multimodal benchmark for wearable stress detection. It contains recordings from 15 subjects who wore a chest-mounted RespiBAN device (capturing EDA, electrocardiogram, electromyogram, respiration, body temperature, and three-axis acceleration at 700 Hz) and a wrist-mounted Empatica E4 during a laboratory protocol that includes baseline rest, an amusement condition, and a stress condition induced by the Trier Social Stress Test. Schmidt et al. reported baseline classification results using hand-crafted statistical features from multiple modalities, achieving up to 93% accuracy with decision trees on combined chest sensor data. Subsequent work has explored deep learning approaches including convolutional neural networks [12], Long ShortTerm Memory (LSTM) networks [13], and attention-based architectures [7] applied directly to raw physiological signals. Recently, an LSTM branch of hybrid model (CNN-LSTM) adopted to process the EDA signals to simulate the temporal fluctuations and sequential dependencies associated with physiological and emotional stress [14]. Most existing approaches frame stress detection as a contemporaneous classification task. Our approach differs by framing it as a forecasting task: given the recent signal, predict characteristics of the near-future signal, and classify from these predictions.

### B. Time Series Foundation Models

Amazon Chronos [9] is a foundation model for time series forecasting that treats continuous time series values as sequences of discrete tokens via quantization into approximately 4096 bins. It is built on the T5 (Text-to-Text Transfer Transformer) architecture [10], an encoder-decoder transformer originally developed for natural language processing that was repurposed for time series by replacing text tokens with quantized numerical values. Chronos models, ranging from 20 million to 710 million parameters, were pretrained on millions of heterogeneous time series using cross-entropy loss over token sequences. The framework achieves strong zero-shot performance across diverse forecasting benchmarks, and has been applied to domains including water quality prediction [15] and energy load forecasting. However, the quantization step may limit precision for signals with subtle amplitude dynamics.

The Tabular Prior-data Fitted Network (TabPFN) [11] is a foundation model for tabular prediction. TabPFN was pretrained on over 100 million synthetic classification and regression tasks and uses a transformer architecture to perform in-context learning: at inference time, it conditions on the training examples provided as context and directly predicts test labels without any gradient-based weight updates. TabPFN has demonstrated strong performance across biomedical and scientific applications [16], and has recently been extended to time series forecasting through lightweight temporal featurization [17].

### C. Tonic/Phasic Decomposition

EDA consists of a slowly varying tonic component (skin conductance level, or SCL) reflecting general arousal, and a fast phasic component consisting of skin conductance responses (SCRs)—transient increases in conductance driven by discrete sympathetic nervous system bursts in response to stimuli. Convex optimization-based decomposition (cvxEDA) [18] provides a principled separation by solving for the smoothest tonic component consistent with a biophysical model of sweat gland dynamics. We employ cvxEDA via the NeuroKit2 library [19] to generate tonic and phasic input channels.

## III. Materials and Methods

### A. Pipeline Overview

The pipeline ran continuously over EDA recordings acquired using the WESAD chest sensor (original 700 Hz) through the following stages: (1) Resampling to 20 Hz followed by a 3 Hz Butterworth lowpass filter; (2) cvxEDA tonic/phasic decomposition; (3) Extraction of overlapping windows, where each window consisted of a 60-second *context window* (1200 samples at 20 Hz) representing the recent past signal that served as input to the forecaster, paired with a *forecast horizon* of 3, 5, or 10 seconds representing the future period over which summary statistics are predicted, using a stride of 2 seconds between consecutive windows; (4) Prediction of three summary statistics describing the forecast horizon: delta mean (the difference between the mean EDA level in the horizon and the current level at the end of the context window), future standard deviation (the variability of the signal within the horizon), and peak delta (the maximum excursion above the current level within the horizon); (5) Binary classification of stress versus baseline using logistic regression with balanced class weights, operating on the three predicted summary statistics as its feature vector.

**Fig. 1.**
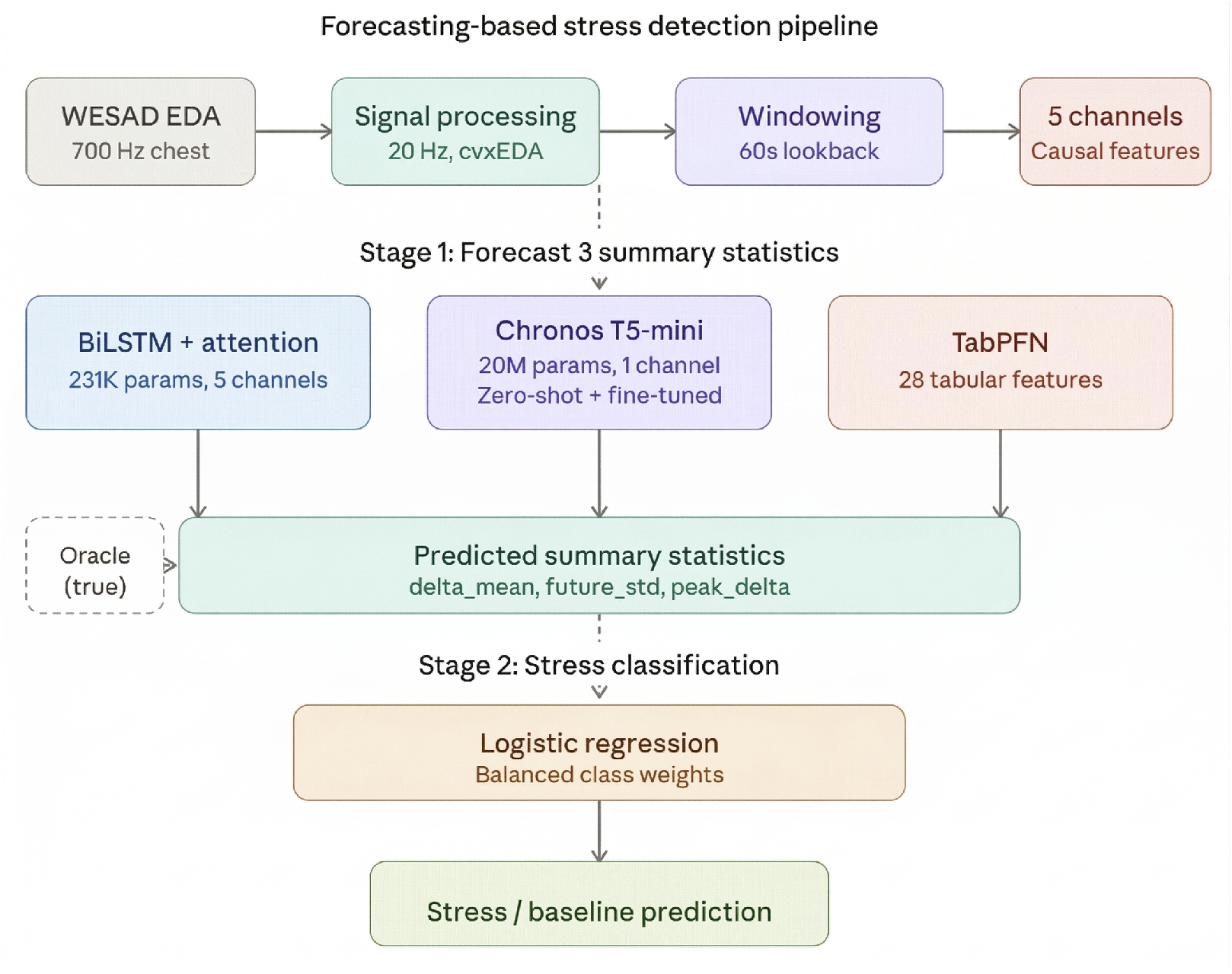
Two-stage pipeline architecture for forecasting-based stress detection. Stage 1 (top): raw 700 Hz EDA from WESAD is downsampled to 20 Hz, decomposed into tonic and phasic components via cvxEDA, and segmented into overlapping 60-second context windows. Three forecasting models—BiLSTM (5 engineered channels), Chronos T5 (univariate input), and TabPFN (28 tabular features)—independently predict three summary statistics of the future signal. Stage 2 (bottom): the predicted statistics serve as a three-dimensional feature vector for a logistic regression classifier that outputs a binary stress/baseline prediction. An oracle branch (dashed) computes the same statistics from the true future signal to establish an upper-bound reference.

All derived input features were computed using strictly causal operations (backward-difference derivatives, trailing-window statistics) to ensure no information from the forecast horizon could leak into the context window.

### B. BiLSTM Architecture

The input to the BiLSTM forecaster had five channels: lowpass-filtered EDA, backward-difference derivative of EDA, trailing 5-second rolling variance of EDA, tonic, and phasic components of EDA from cvxEDA. The architecture was as follows: Bidirectional LSTM layer (128 units per direction) [8], Unidirectional LSTM layer (64 units), layer normalization, learned attention pooling, and two output heads: a regression head predicting the three summary statistics, and an auxiliary trend classification head for regularization. The total number of parameters was approximately 231K, and the training used AdamW with OneCycleLR learning rate scheduling, for up to 80 epochs with early stopping (patience of 15 epochs).

### C. Chronos Configurations

We evaluated chronos-t5-mini (approximately 20 million parameters) in two configurations: (1) *Zero-shot*, where the pretrained model generated probabilistic sample paths from the raw univariate EDA context window and summary statistics were computed from the median trajectory. (2) *Fine-tuned with frozen encoder*, where only the T5 decoder was trained on WESAD EDA segments for 1000 steps at a learning rate of 1 *×*10^*−*3^. Chronos accepts only univariate input and therefore cannot incorporate the derived tonic/phasic channels used by the BiLSTM.

### D. TabPFN Configuration

Rather than processing the raw waveform sequentially, TabPFN received 28 engineered features extracted from each 60-second context window. These included EDA statistics (mean, standard deviation, skewness, kurtosis, range), recent-window statistics (mean and standard deviation of the last 5 and 15 seconds), linear trend slopes computed over the full window and the last 15 seconds, derivative statistics, and tonic/phasic component summaries (mean, slope, and an SCR count proxy based on phasic threshold crossings). Three separate TabPFNRegressor instances predicted each summary statistic independently. TabPFN performed zero-shot inference: it used the training examples as in-context conditioning and predicted without updating any model weights.

### E. Evaluation Protocol

We used subject-independent 5-fold cross-validation, in which the 15 subjects were partitioned into five disjoint groups of three so that no subject appeared in both the training and test sets of any fold. The same fold assignment was used across all models, and for the BiLSTM the early-stopping validation set was drawn from the training subjects rather than the heldout test fold. All reported figures are the mean and standard deviation of the per-fold results.

To evaluate the quality of predicted features, we compared against *oracle features*: the same three summary statistics computed directly from the true future signal rather than from model predictions. Oracle features represent a theoretical upper bound—the best possible features assuming perfect foreknowledge of the future. If predicted features match or exceed oracle performance, the forecasting step preserves or enhances the stress-relevant information in the signal.

We reported AUC (Area Under the Receiver Operating Characteristic Curve) as the primary metric because the test set exhibits substantial class imbalance (approximately 16% stress, 84% baseline). AUC evaluates classification performance across all possible decision thresholds and is therefore robust to imbalanced class distributions, unlike accuracy which can be misleadingly high when a classifier defaults to predicting the majority class.

## IV. Experimental Results

### A. Forecasting Quality

We reported the average of the Pearson correlations (*r*) between the predicted and oracle summary statistics for the three forecast horizons in Table I. Pearson correlation captures the degree of linear agreement between the predicted and true future features, with *r* = 1 being perfect prediction.

**TABLE I.** Average Forecasting Correlation (Pearson *r*) across the three summary statistics (delta mean, future standard deviation, peak delta) for each model and forecast horizon. Higher values indicate more accurate predictions. Bold indicates best per column.

| Pipeline | 3s | 5s | 10s |
| --- | --- | --- | --- |
| BiLSTM EDA | 0.625 | 0.463 | 0.527 |
| Chronos Zero-Shot | 0.529 | 0.500 | 0.416 |
| Chronos FT (Frozen) | 0.548 | 0.513 | 0.465 |
| **TabPFN EDA** | **0.686** | **0.667** | **0.639** |

The TabPFN performed best, achieving the highest average correlation at all horizons and exhibiting stable performance as the forecast horizon increased. The BiLSTM correlated most strongly at a horizon of 3 seconds and decreased at longer horizons. Chronos fine-tuning did not improve over the zeroshot baseline, with the two configurations producing similar correlations.

### B. Stress Classification

We presented the results of stress classification using the predicted features and a logistic regression classifier in Table II.

**TABLE II.** Stress classification performance using forecast features with logistic regression, reported as mean *±*standard deviation of the forecast AUC across the five cross-validation folds for each horizon.

| Pipeline | 3s | 5s | 10s |
| --- | --- | --- | --- |
| BiLSTM EDA | $0.864 \pm 0.139$ | $0.859 \pm 0.134$ | $0.882 \pm 0.110$ |
| TabPFN EDA | $0.883 \pm 0.074$ | $0.876 \pm 0.066$ | $0.863 \pm 0.064$ |
| Chronos Zero-Shot | $0.671 \pm 0.066$ | $0.756 \pm 0.067$ | $0.777 \pm 0.067$ |
| Chronos FT (Frozen) | $0.629 \pm 0.071$ | $0.734 \pm 0.064$ | $0.768 \pm 0.074$ |

The BiLSTM and TabPFN achieved comparable AUC at every horizon; on a paired per-fold basis neither model held a statistically significant advantage over the other, and TabPFN was the more stable of the two, with roughly half the fold-to-fold standard deviation. Both models remained well ahead of the Chronos variants at all horizons. The large standard deviations of the BiLSTM reflect strong inter-subject variability, with one three-subject fold in particular performing well below the others.

Across models, the forecast-based classifiers performed close to their oracle counterparts computed from the true future signal, indicating that the forecasting step preserves most of the stress-relevant information rather than discarding it. For the BiLSTM and TabPFN the forecast AUC was within a few points of the oracle at every horizon, whereas for the Chronos variants the forecast fell well short of the oracle, consistent with a substantial loss of information during tokenized generation.

### C. Fine-Tuning Ablation

Fine-tuning the Chronos decoder did not improve over the zero-shot model; under cross-validation the frozen fine-tuned configuration performed on par with, and at some horizons slightly below, the zero-shot baseline. This indicates that the performance ceiling is architectural rather than a consequence of insufficient training.

## V. Discussion

One significant conclusion is that accuracy of prediction and usefulness of prediction are not necessarily related. TabPFN achieves the highest correlation with oracle, yet under crossvalidation it and the BiLSTM reach comparable downstream classification AUC. This seems to indicate that both a domain-trained sequential model and a foundation tabular model over engineered features are effective routes to EDA based stress discrimination, with TabPFN offering more consistent performance across subjects.

**Fig. 2.**
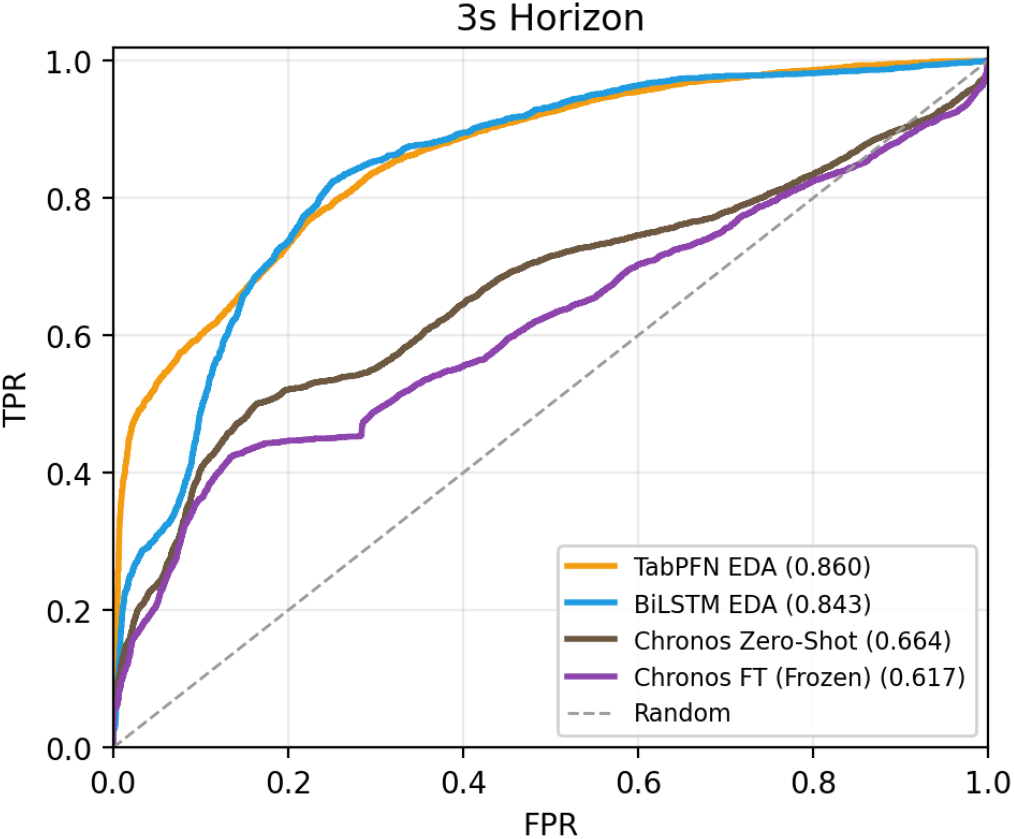
Receiver Operating Characteristic (ROC) curves for stress classification at the 3-second forecast horizon, pooled across cross-validation folds. Each curve plots the true positive rate against the false positive rate across all classification thresholds. TabPFN (AUC = 0.860) and the BiLSTM (AUC = 0.843) separate clearly from both Chronos variants. The diagonal dashed line represents random classification (AUC = 0.50).

**Fig. 3.**
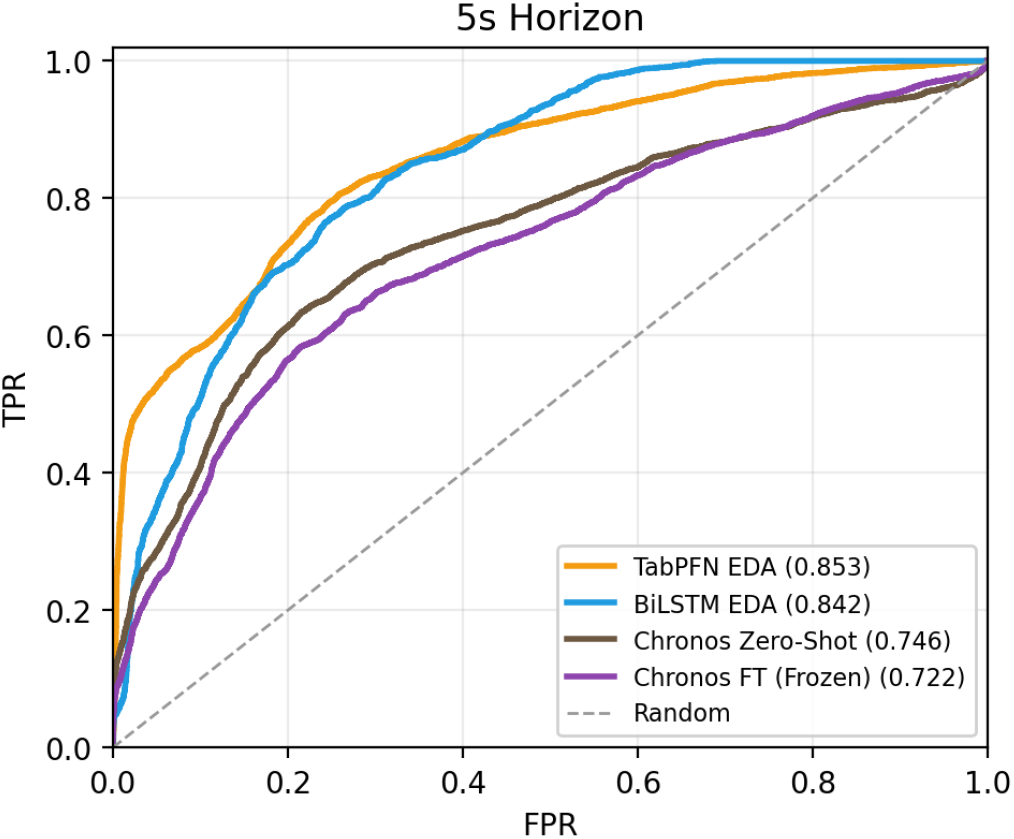
ROC curves for stress classification at the 5-second forecast horizon, pooled across cross-validation folds. TabPFN (AUC = 0.853) and the BiLSTM (AUC = 0.842) are closely matched. Chronos fine-tuning does not improve over the zeroshot baseline, and both Chronos variants remain substantially below the BiLSTM and TabPFN.

**Fig. 4.**
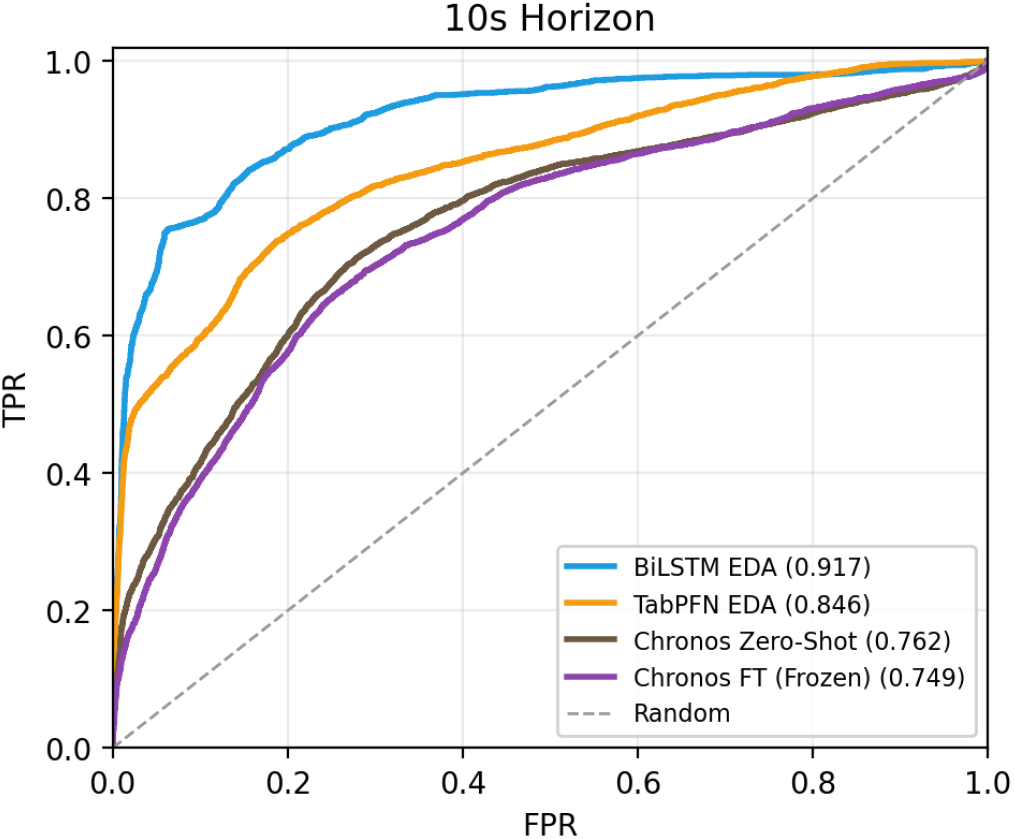
ROC curves for stress classification at the 10-second forecast horizon, pooled across cross-validation folds. The BiLSTM reaches the highest AUC (0.917) at this horizon, with TabPFN (0.846) close behind, while the two Chronos variants cluster together near AUC 0.75.

These findings also have implications beyond stress detection, particularly for human-centric digital twins that require individualized, temporally resolved indicators of behavior. Inter-individual variation in physiological stress reactivity can reveal behavioral signatures relevant to domains where traditional surveillance tools lack sensitivity, such as nutrition and nutrition insecurity. Because nutritional vulnerability is shaped by dynamic interactions between stress, coping behaviors, and environmental constraints, forecasting-based physiological modeling may provide a scalable pathway for capturing individualized behavioral patterns that are otherwise difficult to observe. This suggests that physiological digital twins incorporating predictive stress features could enhance precision monitoring in contexts where static or self-reported measures are insufficient.

Chronos results also suggest that tokenization-based time series models are poorly suited for physiological signal tasks. 20M model with pre-training on millions of time-series is unable to perform even comparable to the BiLSTM which has 231K parameters and is trained on this domain specific data. This is not an adaptation problem since with added training, Chronos actually performs similar or even worse, suggesting that losing the information that the amplitude values encode during the tokenization process removes important features for EDA based stress detection.

TabPFN is an interesting middle ground as it incorporates both tonics and phasics like the BiLSTM, but without capturing the temporal dependency like Chronos. The fact that it achieves consistent performance regardless of horizon (0.863-0.883) seems to suggest that the summary statistics regarding the physiological signal capture the vast majority of the stress-predictive information available and leverageable by the sequential model.

Our findings reinforce the architectural choices seen in existing hybrid CNN-LSTM and Temporal Convolutional Autoencoder (TCAE) frameworks designed to capture complex temporal fluctuations [14], [5]. On the strongest folds the BiLSTM approaches the AUC-ROC of 0.963 achieved by the TCAE model in [5], though its cross-validated mean (0.859–0.882) is lower and more variable across subjects.

## Limitations

Although we used subject-independent crossvalidation, the dataset contains only 15 subjects, so the per-fold estimates carry wide uncertainty and the strong intersubject variability we observe would be better characterized on larger and more diverse cohorts. The cvxEDA decomposition that was used to derive tonics and phasics is an offline procedure requiring optimization on the entire signal; ideally causal methods are needed for real-time scenarios. In addition, there is a subtle asymmetry, the BiLSTM uses 5 input channels whereas Chronos uses 1 univariate signal; this is an architectural decision of the pre-trained Chronos and not something designed as part of the experiment. Furthermore, we only focus on EDA signals and it is natural to move toward incorporating respiration and heart rate signals.

## VI. Conclusion

We presented a forecasting-based approach to EDA stress detection comparing domain-trained and foundation model architectures under subject-independent cross-validation. The domain-trained BiLSTM (0.859–0.882 AUC) and the TabPFN foundation model (0.863–0.883 AUC) achieved comparable performance, with TabPFN being the more stable across subjects, and both substantially outperformed Chronos (0.629– 0.777). A fine-tuning ablation confirms that the performance ceiling of Chronos is architectural rather than training-limited. These findings suggest that for physiological time series, domain-informed models—whether sequential or featurebased—remain more effective than general-purpose tokenized foundation models.

## Notes

### Competing Interest Statement

The authors have declared no competing interest.

### Summary of Updates

This version replaces the single fixed train/test split used in the previous version with subject-independent 5-fold cross-validation, in which the 15 subjects are partitioned into five disjoint groups of three so that no subject appears in both the training and test sets of any fold. All reported results, tables, and figures have been updated accordingly, and now report the mean and standard deviation across folds rather than single-split values. Under cross-validation, the BiLSTM and TabPFN achieve comparable classification performance (mean AUC 0.859 to 0.882 and 0.863 to 0.883 respectively), with TabPFN being the more stable across subjects, and both remain well ahead of the Chronos variants (0.629 to 0.777). The claims in the abstract, results, discussion, and conclusion have been adjusted to reflect this, replacing the earlier statement that the BiLSTM achieved the highest performance. The Chronos unfrozen fine-tuning configuration has been removed, and the analysis now compares zero-shot Chronos against a single frozen-encoder fine-tuned configuration. The fine-tuning ablation has been updated to note that fine-tuning does not improve over zero-shot under cross-validation, which reinforces the conclusion that the performance ceiling of Chronos is architectural rather than training-limited. The oracle versus forecast comparison table has been removed, as the advantage of forecast features over oracle features largely disappears under cross-validation. The corresponding text now states that forecast-based classifiers perform close to their oracle counterparts rather than exceeding them. Table I (forecasting correlation) and Table II (classification performance) have been updated to cross-validated values, and the ROC curve figures have been regenerated from the pooled cross-validation predictions. Additional digital twin references have been added and the related discussion expanded. The evaluation protocol and limitations sections have been revised to describe the cross-validation procedure and to note the wide per-fold uncertainty given the small number of subjects.

